# AntiOxiPred: Predicting Antioxidant Peptides using Machine Learning Techniques and BLAST

**DOI:** 10.64898/2026.10.21.760867

**Authors:** Adrita Kundu, Nisha Bajiya, Gajendra P.S. Raghava

**Affiliations:** Department of Biological Sciences, Indian Institute of Science Education and Research, Mohali, Punjab, India; Department of Computational Biology, Indraprastha Institute of Information Technology, New Delhi, India

**Keywords:** Antioxidant peptides, Machine learning, Ensemble approach, BLAST, Reactive oxygen species

## Abstract

Antioxidant peptides (AOPs) offer significant therapeutic potential in counteracting oxidative damage associated with chronic diseases. As AOPs gain market traction in cosmeceuticals, functional foods, and clinical trials for disease intervention, efficient computational methods are essential to accelerate their discovery. In this study, a systematic attempt has been made to develop a highly accurate and reliable method for predicting AOPs from their primary sequences. The dataset comprises 1,511 experimentally validated AOPs and 1,511 randomly generated non-AOPs, partitioned into 2,417 sequences for training and 606 for testing. First, we developed machine learning-based models using one-hot encoded (OHE) representations, sequence-derived compositional features, protein language model (PLM) embeddings, and feature fusion. Our CatBoost model, developed using selected fused features, achieved the best performance, with an AUC of 0.945 and an MCC of 0.788 on the independent testing dataset. An ensemble model combining CatBoost with BLAST achieved a peak AUC of 0.956 on the same testing dataset. Furthermore, our method outperformed existing state-of-the-art predictors when benchmarked on two external validation datasets comprising 150 peptides (75 AOPs and 75 non-AOPs) and 248 peptides (124 AOPs and 124 non-AOPs). AntiOxiPred (https://webs.iiitd.edu.in/raghava/antioxipred/) is freely available as an interactive web server, a standalone package, and a pip-installable tool to support the scientific community

## 1. Introduction

Reactive oxygen species (ROS) are highly reactive oxygen-containing free radicals which are generated naturally as by-products of aerobic metabolism. These free radicals are continuously produced in biological systems through both endogenous and exogenous pathways.[1][2][3][4][5] Under physiological conditions, ROS act as essential secondary messengers in multiple signalling pathways and participate in the regulation of cell proliferation, differentiation, apoptosis, immune defence, wound healing, and adaptation to environmental stress.[6][7][8][9][10] Antioxidants play a crucial role in scavenging excess ROS and maintaining intracellular redox balance at levels compatible with normal cellular functioning. These defence systems prevent the uncontrolled accumulation of ROS, thereby protecting biomolecules from oxidative injury and preserving cellular homeostasis.[11][12][13] An imbalance between ROS generation and antioxidant defence systems results in oxidative stress, during which excess ROS can induce oxidative damage to critical biomolecules, including lipids, proteins, carbohydrates, and nucleic acids. Persistent oxidative stress can consequently lead to the pathogenesis of various chronic diseases, as shown in Figure 1.[12][14][15][16] Synthetic antioxidants such as butylated hydroxyanisole (BHA), butylated hydroxytoluene (BHT), tert-butylhydroquinone (TBHQ), and propyl gallate were extensively introduced into the food, cosmetic, and pharmaceutical industries to minimize oxidative deterioration because they have strong radical scavenging activity.[17][18] But extensive studies have provided toxicological evidence regarding the long-term safety of several synthetic antioxidants.[19][20] Growing concerns regarding the toxicity and long-term safety of synthetic antioxidants have accelerated interest in naturally derived antioxidant compounds, particularly antioxidant peptides (AOPs), which have emerged as promising candidates for applications in food preservation, nutraceuticals, pharmaceuticals, and the management of oxidative stress-associated diseases.[21][22][23] They have been identified from a wide variety of natural sources, including milk, egg, soybean, wheat germ, rice bran, fish, marine organisms, algae, edible insects, fermented foods, and microbial protein hydrolysates, and more than 1000 AOPs have been reported.[24] However, conventional experimental approaches for AOP discovery, including isolation, purification, and activity screening, are labor-intensive, time-consuming, and economically demanding, thereby necessitating the development of efficient computational strategies for rapid AOP prediction and screening.[25]

**Figure 1:**
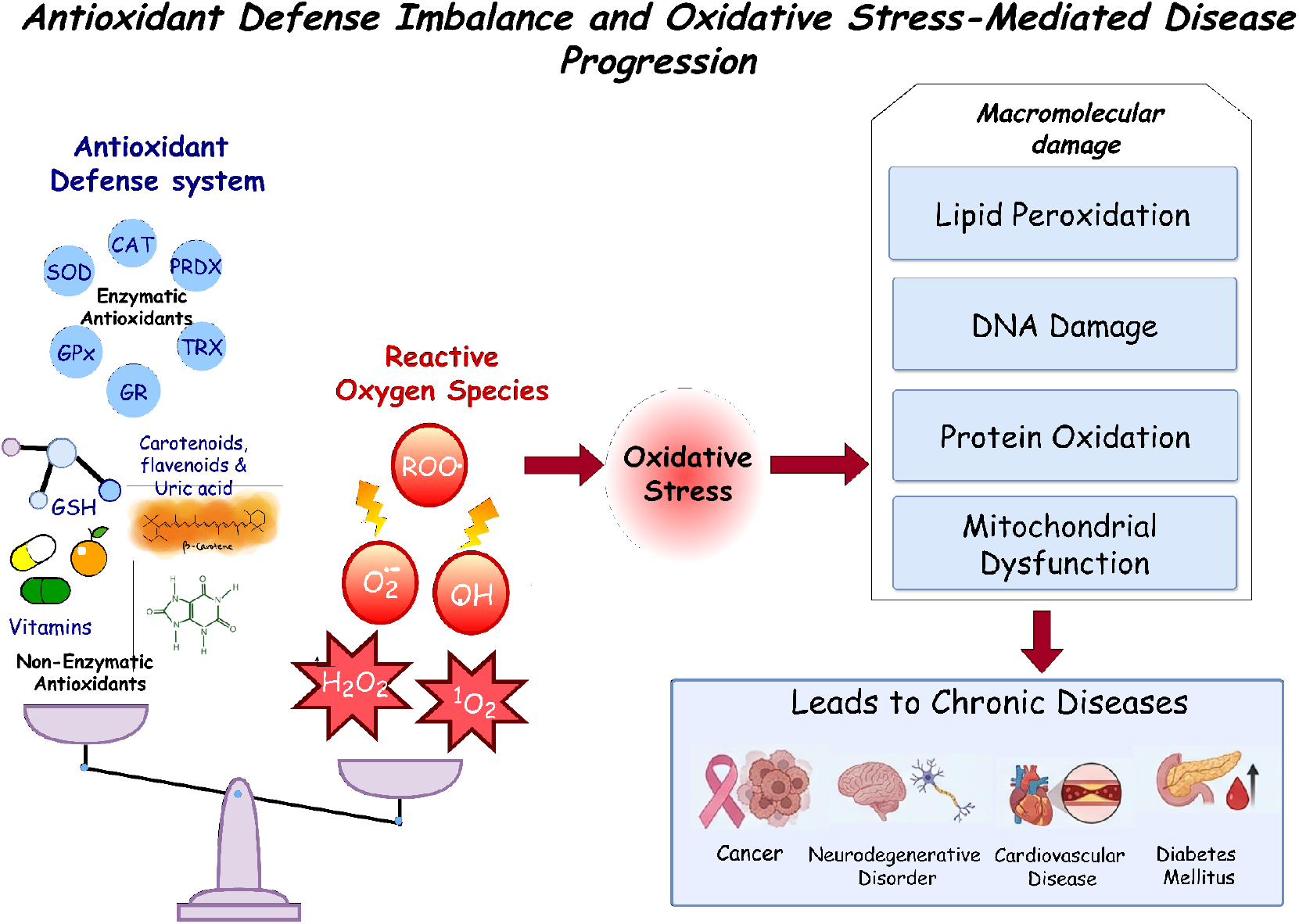
The imbalance between antioxidant defence systems and ROS leads to oxidative stress, macromolecular damage, and chronic disease progression.

One of the earliest deep learning-based predictors and the first online server for antioxidant prediction, AnOxPePred, was introduced in 2020 and uses convolutional neural networks (CNNs) trained on experimentally validated antioxidant and non-antioxidant peptides.[26] The model categorized peptides into free radical scavenging and metal-chelating classes and demonstrated improved performance over traditional k-NN based approaches. Subsequently, AnOxPP was introduced in 2023 using a bidirectional long short-term memory (Bi-LSTM)-based quantitative structure-activity relationship framework integrated with interpretable amino acid descriptors. The model achieved accuracies of 0.967 and 0.819 on two independent datasets; however, the considerable discrepancy between these performances suggested limitations in robustness and generalization across diverse peptide datasets.[27] To further improve robustness and feature representation, several machine learning (ML)-based predictors were introduced; iAnOxPep utilized ensemble learning with multiple sequence-derived descriptors to improve prediction stability across the benchmark dataset.[28] Similarly, a BiLSTM-based model, AOPP [29] and AOPxSVM.[30] which integrated SVM with BLOSUM62 and CTD to enhance multidimensional sequence characterization showed accuracies exceeding 90% with improved independent dataset performance compared with earlier predictors, demonstrating superior classification robustness across benchmark datasets. More recently, AOP-DRL employed protein language model (PLM) embeddings integrated with Text-CNN architectures to capture nonlinear sequence relationships and variable peptide lengths more efficiently.[31]

Although computational methods for predicting AOPs have improved considerably, many models still perform inconsistently on independent datasets. This limits their ability to generalize to unseen data and reduces their practical application. To address these problems, we compared 10 ML algorithms across multiple feature representations and selected the best-performing model based on its average ranking across evaluation metrics, providing a consistent basis for comparison. An ensemble model integrating our best-performing CatBoost model with BLAST alignment was also tested to assess whether it yielded improved performance. Finally, we developed a web server and a standalone software tool, AntiOxiPred, for the effective prediction of antioxidant peptides.

## 2. Materials and Methods

### 2.1. Collection and Preprocessing of the Dataset

The AOP dataset used for model development in the present study was obtained from the previously published AOPP framework.[29] The dataset was curated from multiple public repositories, including BIOPEP-UWM,[32] AODB,[33] PlantPepDB,[34] and FermFooDb,[35] resulting in 1511 experimentally validated AOPs ranging in length from 2 to 20 amino acids. An equal number of negative peptides were generated in the original study, and the dataset was then reduced for redundancy using CD-HIT [36] at a 90% similarity threshold. The final dataset consisted of 3022 sequences, which were split into training and test sets in an 80:20 ratio, yielding 2417 and 606 sequences, respectively.

Additionally, an independent validation dataset 1 consisting of 75 experimentally validated AOPs reported between 2022 and 2023 was adopted from the AOPP study. A second independent dataset of 125 experimentally validated AOPs reported between 2022 and 2026 was manually curated in the present study. All peptides in the second independent dataset 2 were verified to be unique relative to the training, test, and first independent datasets. The test dataset, comprising 606 sequences, was used to develop the model, whereas the independent datasets were subsequently used to benchmark against previously reported prediction models.

### 2.2. Amino Acid Composition and Length Distribution Analysis

Amino Acid Composition (AAC) analysis was performed to determine the relative abundance of each amino acid within peptide sequences. AAC represents the normalized frequency of occurrence of individual amino acid residues and was calculated using the following equation 1:

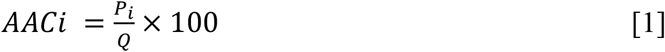

Here, *AACi* denotes the composition of amino acid *i*, *Pi* represents the number of residues of type *i*, and Q corresponds to the total peptide length. Additionally, peptide length distribution analysis was conducted to characterize the sequence length variability within the positive dataset.

### 2.3. Positional Preference Analysis

To identify position-specific residue preferences for antioxidant and non-antioxidant peptides, a Two-Sample Logo analysis (TSL) [37] was performed. As most of the peptides in our dataset comprised 3 to 5 residues, we considered the first 3 N-terminal and last 3 C-terminal residues for the analysis. This allowed us to compare similar positions for peptides of different lengths. To prepare the TSL, sequences from the positive and negative sets were used as input, enabling the identification of enriched or depleted residues at specific positions in the two groups.

### 2.4. Feature Extraction

#### 2.4.1. Composition-based features

To capture the features of the positive and negative datasets, a diverse set of composition and sequence-based descriptors was generated using Pfeature.[38] We extracted Compositional features such as AAC, Dipeptide Composition (DPC), Atom Composition (ATC), and Bond Composition (BTC), which describe amino acid composition, local sequence patterns, and the chemical constitution of peptides. Physicochemical Properties (PCP), Amino Acid Index (AAI), Distance Distribution of Residues (DDR), Repetitive Residue Information (RRI), and Composition-enhanced Transition Distribution (CeTD) were also extracted to characterize residue abundance, positional distribution, repetition patterns, and physicochemical variation within peptide sequences. To capture higher-order sequence information, we generated Sequence Order Coupling Number (SOCN), Quasi-Sequence Order (QSO), Property Repeats (PRI), Shannon entropy-based descriptors (SER, SEP, and SPC), Pseudo Amino Acid Composition (PAAC), and Amphiphilic Pseudo Amino Acid Composition (APAAC).

#### 2.4.2. One-Hot Encoding representation

One-Hot Encoding (OHE) encodes each amino acid in a peptide sequence as a binary vector. In this approach, each standard amino acid is assigned a unique vector of length 20, with one position set to 1 (indicating the presence of that amino acid) and all other positions set to 0. For example, if alanine (A) is assigned the first position, it is represented as (1, 0, 0,…,0). For a peptide of length L, we produce a feature matrix of size L × 20, which is typically flattened into a vector of length 20L for model input. Since peptide sequences vary in length, all sequences were padded to a uniform length (20 residues in this study) using a dummy residue (X). Positions corresponding to padding are encoded as all zeros, ensuring that no artificial signal is introduced.

#### 2.4.3. PLM Embeddings

Pre-trained PLM embeddings capture contextual and evolutionary information from peptide sequences. Embeddings were generated using ProtBERT,[39] BioBERT,[40] and multiple ESM2 variants (8M, 35M, and 150M parameters).[41] For each model, peptide sequences were provided as input in raw amino-acid format rather than as numerical features. In transformer-based architectures such as ProtBERT and ESM2, sequences were tokenized at the residue level and passed through multiple attention layers to generate context-aware embeddings, where each residue representation is influenced by its neighbouring sequence context.

ProtBERT embeddings were derived from the ProtTrans framework, while BioBERT, originally developed for a huge number of biomedical texts, follows the BERT architecture and has been adapted for biological sequence representations. ESM2 models, trained on large-scale protein sequence datasets, provide rich evolutionary representations that enhance downstream predictive performance. Pretrained weights were used without further fine-tuning to ensure consistency and reproducibility.

### 2.5. Classification Models

To evaluate the predictive performance of the extracted features, multiple ML classifiers representing diverse learning paradigms were employed. These included ensemble-based methods such as Random Forest (RF), Decision Tree (DT), Categorical Boosting (CatBoost), Extreme Gradient Boosting (XGBoost), and Light Gradient Boosting Machine (LGBM), kernel-and linear-based models including Support Vector Machine (SVM) and Logistic Regression (LR), neural network-based Multi-Layer Perceptron (MLP), instance-based K-Nearest Neighbors (KNN), and the probabilistic classifier Gaussian Naive Bayes (GNB). The hyperparameters of the models were optimized using GridSearchCV from the scikit-learn package. The ideal combination of hyperparameters for each classifier was selected based on the highest area under the ROC curve (AUC) metric.

### 2.6. Feature Fusion and Selection

To add complementary sequence information, feature fusion was performed by concatenating standardized feature vectors to curate high-dimensional representations for each peptide. After this, keeping in mind that feature fusion can also introduce redundancy, we performed feature selection to reduce and retain the optimal set of attributes. Feature selection was performed using a Support Vector Classifier with L1 regularization (SVC-L1). This model uses a penalty to shrink the weights of less important features down to zero.

### 2.7. Model Evaluation and Performance Metrics

To optimize the classification models and minimize overfitting, a 5-fold cross-validation technique was used. In this approach, the dataset is randomly divided into five equal sets; in each iteration, four sets are used for training and the remaining set for validation. This process is repeated 5 times, with each set used exactly once as a validation set, and the final performance is reported as the mean across all folds. The performance of the model is evaluated based on multiple metrics from equations 2 to 6, with the Area Under the Receiver Operating Characteristic Curve (AUC) used to determine the overall discriminative power of the model, complemented by Accuracy (ACC), Sensitivity (Sn), Specificity (Sp), and Matthews Correlation Coefficient (MCC).

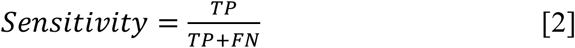

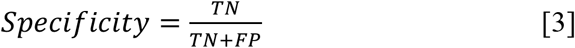

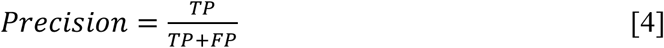

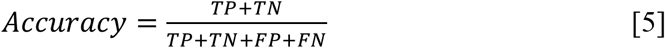

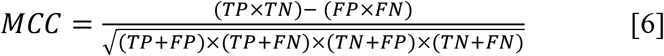

### 2.8. Similarity Search Method - BLAST

Similarity-based annotation was performed using BLASTp-short,[42] which is designed to handle short peptide sequences, to classify peptides as antioxidant or non-antioxidant based on sequence similarity. The custom-made BLAST database was prepared from the training set using *makeblastdb*. The query sequences were then compared against the created database with different E-value thresholds. For training data evaluation, the search results were classified by the first non-self-hit’s score, whereas for the validation set, the closest database entry was selected as a hit based on the top result’s score. The classification was based on the classification of the closest homologs, since similar sequences are likely to be functionally similar, as demonstrated in proteomics and peptides.[43][44]

### 2.9. Ensemble Approach

We also hypothesized that the model’s predictive performance can increase using an ensemble strategy. In this ensemble approach, we implemented a weighted scoring method that combines two techniques to calculate the score: a similarity-based approach using BLAST and a final classification model. For each query, the top hit was considered for classification. Based on the nature of the hit, a score was assigned as follows: if the query sequence aligned with an AOP, a score of +0.5 was assigned; if it aligned with a non-AOP, a score of-0.5 was assigned; and if no significant hit was obtained, a score of 0 was assigned. This scoring scheme allows the incorporation of homology-based evidence into the prediction pipeline, in which sequence similarity quantitatively contributes to the final decision.[43]

### 2.10. Web server implementation

An interactive web server, AntiOxiPred, was developed using HTML5, CSS3, JavaScript (ES6), PHP (version 8.2.12), and Python (version 3.13.5) to enable rapid AOP prediction. The server provides a user-friendly interface for uploading sequences and obtaining predictions, with downloadable results. To facilitate accessibility and reproducibility, a standalone Python implementation of AntiOxiPred and a pip-installable package were made available.

### 2.11. Statistical Analysis

Statistical analyses were conducted to understand feature-level differences and classifier performance. For AAC comparisons, Welch’s t-test was applied using the *scipy.stats module*, as it does not assume equal variances. To correct for multiple comparisons across amino acids, Benjamini-Hochberg false discovery rate (FDR) correction was applied using *statsmodels.stats.multitest.*[45] For classifier comparison, a Friedman test [46] was used to detect overall differences among models, followed by a Nemenyi post hoc test for pairwise comparisons. This analysis was performed using GraphPad Prism (version 11.0.0) and visualized through a critical difference (CD) diagram, where overlapping ranks indicate no significant difference.

## 3. Results

The complete workflow of the study is demonstrated in Figure 2.

**Figure 2:**
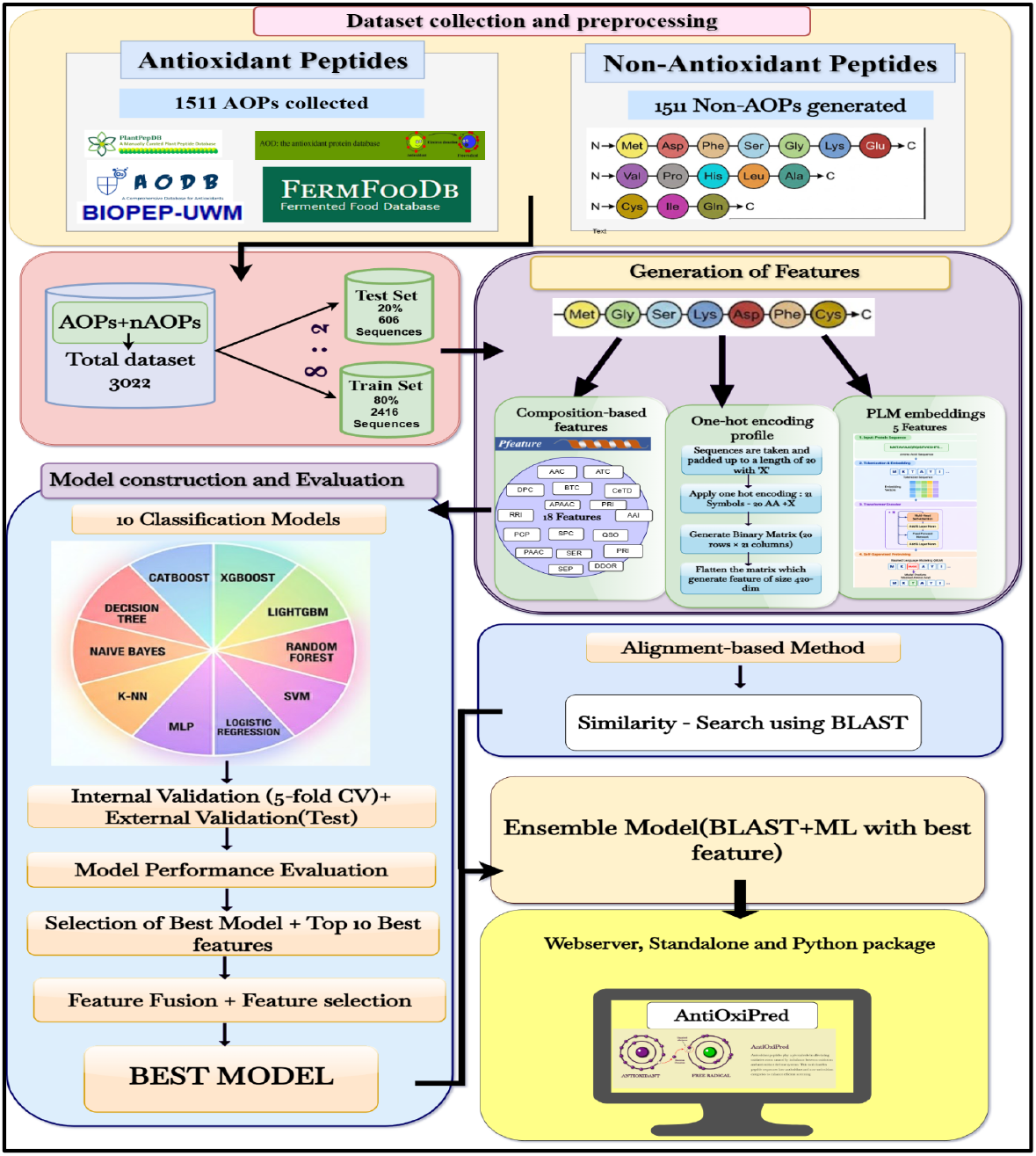
The architecture of AntiOxiPred.

### 3.1. Analysis of AOPs

#### 3.1.1. Peptide Length Distribution Analysis

The peptide length distribution for the AOP dataset is demonstrated in Figure 3 below. The length varied from 2 to 20 amino acids, out of which the peptide length of 3 amino acids was the most common. Another peak was observed for 5-residue peptides. The number of peptides gradually decreased as the length increased. Longer chains were represented rarely, and those containing more than 15 amino acid residues were barely present. As seen in the graph, the tendency reflects the common occurrence of short-chain peptides, a characteristic commonly associated with bioactive peptides and their favorable biological properties.

**Figure 3:**
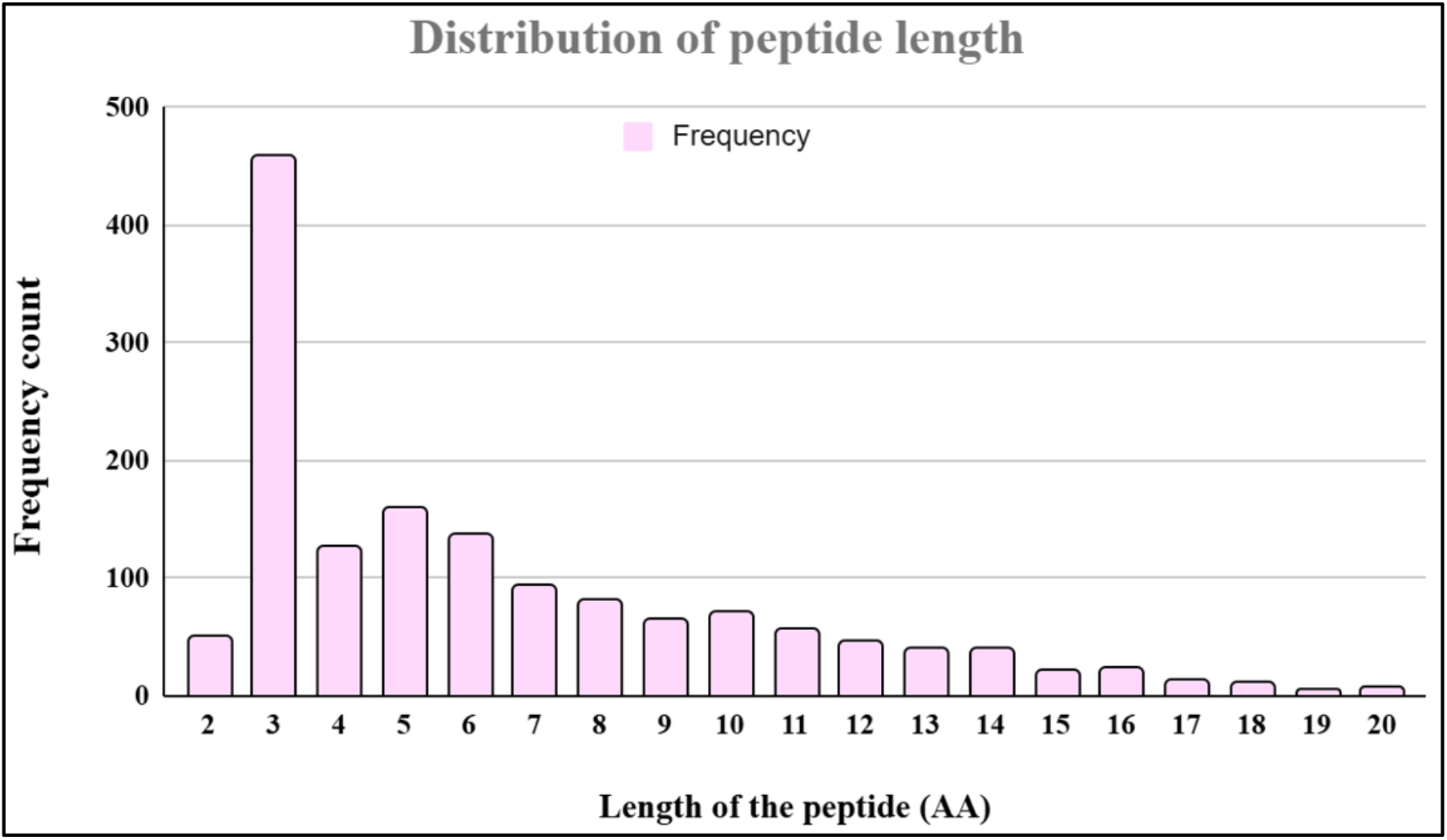
Histogram plot showing the frequency distribution of the main positive dataset.

#### 3.1.2. AAC Analysis

The investigation into the AAC of AOPs and non-AOPs in Figure 4 revealed distinct patterns that underscore their functional differences. Also, the corresponding mean composition values and standard deviations for all amino acids are provided in the Supplementary Table S1. Several residues, including Leucine (L), Proline (P), Tyrosine (Y), Alanine (A), Glutamic acid (E), Glycine (G), Histidine (H), and Valine (V), were significantly enriched (*p* < 0.001) in AOPs. Notably, Y and H are known to contribute to radical scavenging and metal-chelating activities. At the same time, hydrophobic residues such as L, V, A, and Y may enhance interactions with lipid-associated radicals. In contrast, Cysteine (C), Methionine (M), Asparagine (N), Glutamine (Q), Arginine (R), Threonine (T), Tryptophan (W), Phenylalanine (F), and Isoleucine (I) were more abundant in non-AOPs. Interestingly, despite the reported antioxidant roles of residues such as C, W, and M, their enrichment in non-AOPs suggests that antioxidant activity depends on overall sequence composition and residue arrangement rather than on the presence of individual amino acids. Aspartic acid (D) and Lysine (K) showed no significant differences between the two classes, indicating that these residues may have a neutral or less discriminatory role in distinguishing AOPs from non-AOPs.

**Figure 4:**
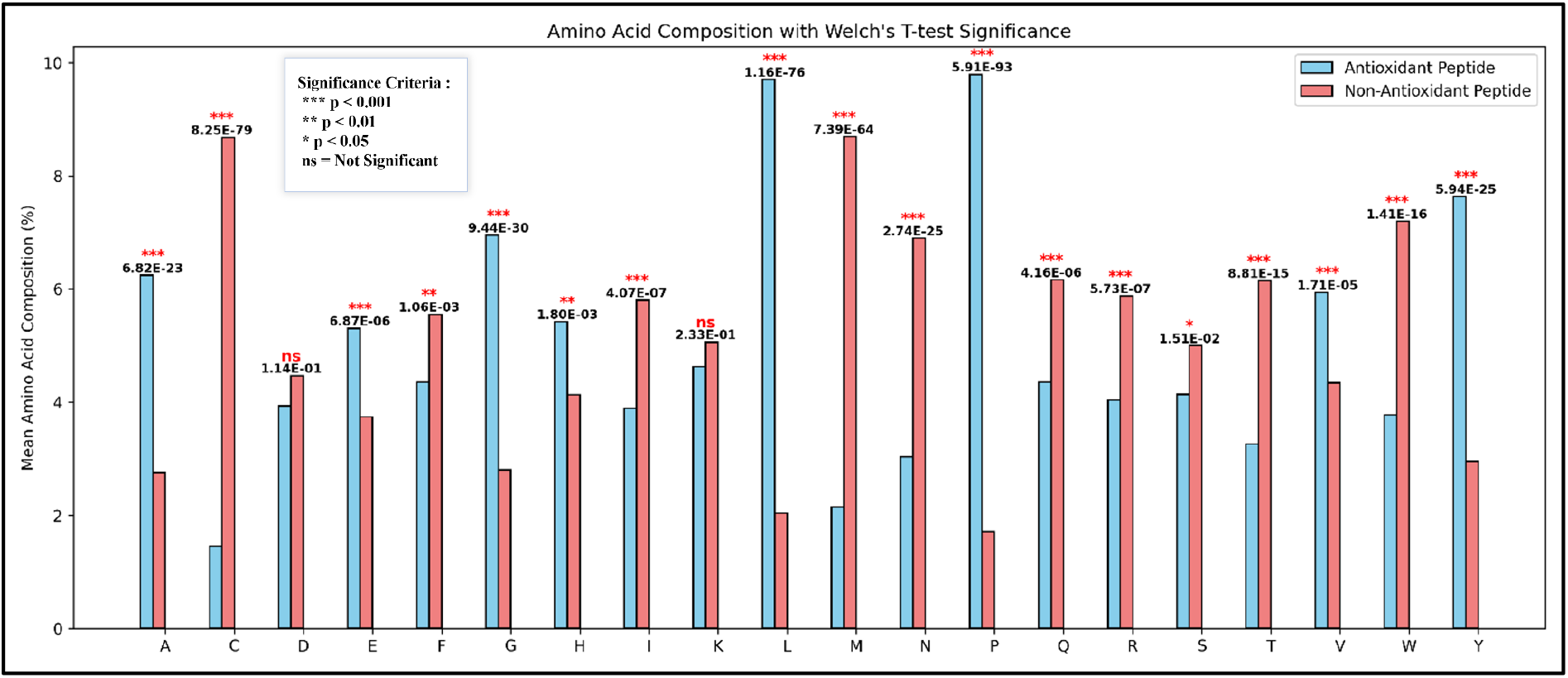
Bar plots of the percentage average AAC comparison between AOPs and non-AOPs.

#### 3.1.3. Positional Residue Analysis

To understand the residue preference at a specific position associated with antioxidant activity, a position-specific analysis was performed using TSL. In agreement with the compositional analysis, the TSL in Figure 5 showed that the AOPs exhibit a visible enrichment of L, P, Y, and A amino acids in the N- and C-terminal regions. The abundance of hydrophobic residues such as leucine and tyrosine may facilitate interactions with lipid-associated radicals, while proline likely contributes to structural stability. In contrast, non-AOPs were abundant in residues such as M, C, and N.

**Figure 5:**
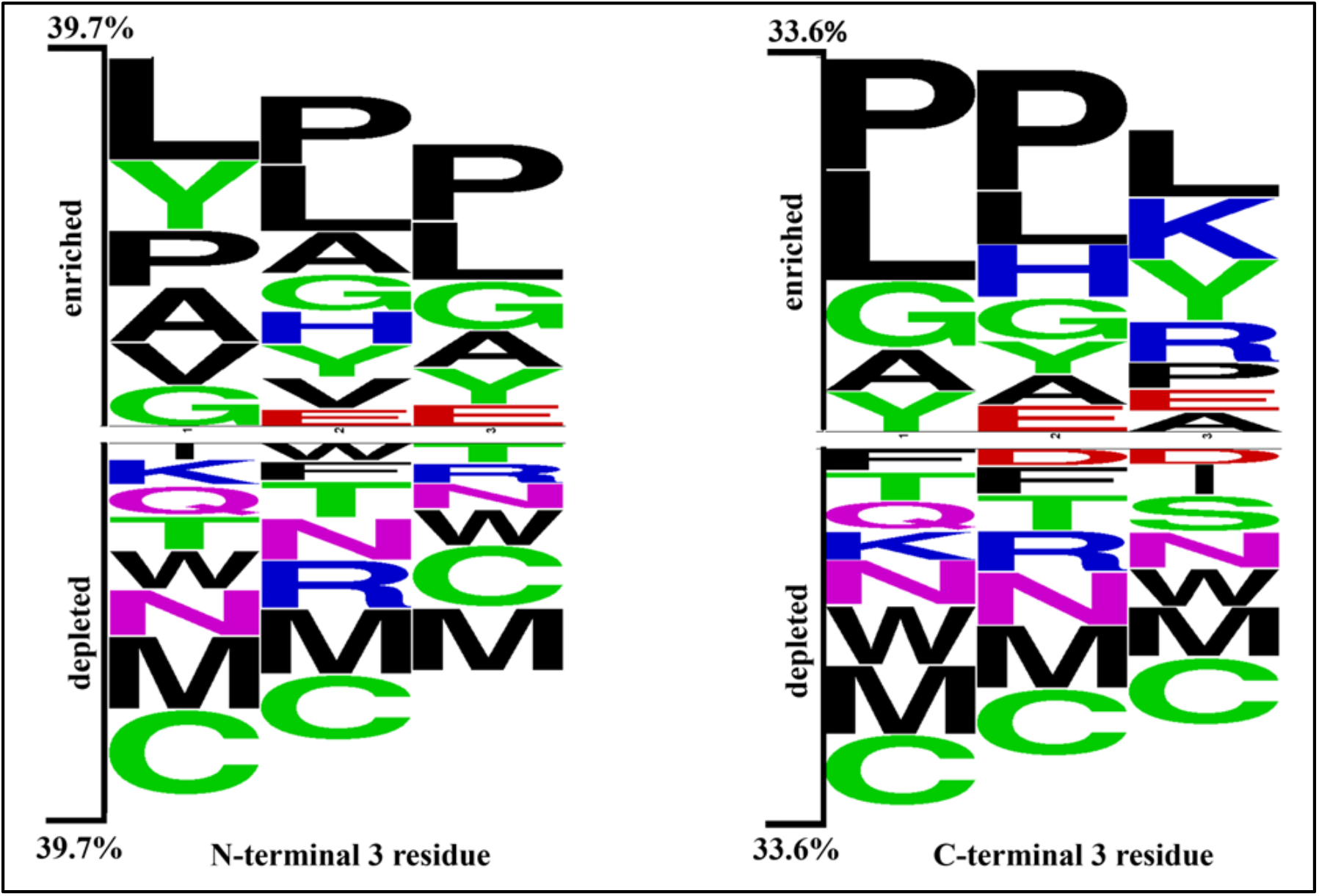
Three residue preferences in AOPs and non-AOPs at N- & C-terminal.

### 3.2. Classification Models

#### 3.2.1. Performance of Compositional and OHE features

To identify the most informative feature representations, all 18 sequence feature descriptors were evaluated using ten different ML algorithms. The classifier’s performance on the top compositional feature and OHE is shown in Tables 1 and 2, and the complete results are provided in Supplementary Tables S2 and S3, respectively. On the independent testing set, QSO with XGB achieved the highest overall performance, with AUC and MCC of 0.939 and 0.775, respectively, while on OHE, CatBoost showed the highest AUC of 0.937 and MCC of 0.773. The strong and consistent performance of these descriptors suggests that they capture complementary sequence characteristics associated with antioxidant activity and were therefore selected for subsequent feature fusion and optimization analyses.

**Table 1:** The performance of the best-performing machine learning-based models on the top compositional features.

| Models | Features | Training set |  |  |  |  | Testing set |  |  |  |  |
| --- | --- | --- | --- | --- | --- | --- | --- | --- | --- | --- | --- |
|  |  | Sn | Sp | ACC | AUC | MCC | Sn | Sp | ACC | AUC | MCC |
| XGB | QSO | 0.853 | 0.897 | 0.875 | 0.943 | 0.751 | 0.845 | 0.927 | 0.886 | 0.939 | 0.775 |
| XGB | PAAC | 0.866 | 0.901 | 0.884 | 0.945 | 0.768 | 0.835 | 0.904 | 0.87 | 0.937 | 0.741 |
| CatBoost | AAC | 0.851 | 0.907 | 0.879 | 0.942 | 0.759 | 0.842 | 0.914 | 0.878 | 0.935 | 0.758 |
| CatBoost | APAAC | 0.855 | 0.903 | 0.879 | 0.943 | 0.758 | 0.838 | 0.917 | 0.878 | 0.933 | 0.758 |
| CatBoost | RRI | 0.831 | 0.912 | 0.871 | 0.938 | 0.745 | 0.848 | 0.901 | 0.875 | 0.932 | 0.75 |
| CatBoost | SER | 0.852 | 0.895 | 0.874 | 0.942 | 0.748 | 0.828 | 0.931 | 0.88 | 0.932 | 0.763 |
| CatBoost | AAI | 0.858 | 0.896 | 0.877 | 0.939 | 0.755 | 0.818 | 0.927 | 0.873 | 0.93 | 0.75 |
| XGB | DDOR | 0.85 | 0.824 | 0.875 | 0.939 | 0.752 | 0.848 | 0.822 | 0.875 | 0.929 | 0.75 |
| MLP | PCP | 0.868 | 0.865 | 0.867 | 0.93 | 0.734 | 0.822 | 0.931 | 0.876 | 0.926 | 0.757 |

#Sn: Sensitivity, Sp: Specificity, ACC: Accuracy, AUC: Area under ROC curve, MCC: Matthews Correlation Coefficient

**Table 2:** The performance of machine learning-based models developed using one-hot encoded features.

| Models | Training set |  |  |  |  | Testing set |  |  |  |  |
| --- | --- | --- | --- | --- | --- | --- | --- | --- | --- | --- |
|  | Sn | Sp | ACC | AUC | MCC | Sn | Sp | ACC | AUC | MCC |
| **CatBoos**<br>**t** | **0.842** | **0.917** | **0.88** | **0.937** | **0.761** | **0.832** | **0.937** | **0.884** | **0.937** | **0.773** |
| XGB | 0.823 | 0.91 | 0.867 | 0.932 | 0.736 | 0.822 | 0.931 | 0.876 | 0.93 | 0.757 |
| LGBM | 0.808 | 0.93 | 0.869 | 0.927 | 0.743 | 0.785 | 0.95 | 0.868 | 0.922 | 0.746 |
| RF | 0.779 | 0.948 | 0.863 | 0.928 | 0.738 | 0.759 | 0.957 | 0.858 | 0.92 | 0.731 |
| SVM | 0.803 | 0.917 | 0.86 | 0.925 | 0.726 | 0.815 | 0.927 | 0.871 | 0.92 | 0.747 |
| LR | 0.773 | 0.916 | 0.844 | 0.913 | 0.697 | 0.749 | 0.924 | 0.837 | 0.909 | 0.684 |
| MLP | 0.857 | 0.881 | 0.869 | 0.928 | 0.738 | 0.832 | 0.875 | 0.853 | 0.906 | 0.707 |
| KNN | 0.6 | 0.38 | 0.49 | 0.912 | 0 | 0.769 | 0.941 | 0.855 | 0.911 | 0.72 |
| NB | 0.803 | 0.829 | 0.816 | 0.889 | 0.633 | 0.822 | 0.835 | 0.828 | 0.89 | 0.657 |
| DT | 0.415 | 0.933 | 0.674 | 0.688 | 0.408 | 0.442 | 0.934 | 0.688 | 0.696 | 0.432 |

#Sn: Sensitivity, Sp: Specificity, ACC: Accuracy, AUC: Area under ROC curve, MCC: Matthews Correlation Coefficient

#### 3.2.2. Performance of Protein Learning Embeddings

Five embedding representations were assessed using 10 ML algorithms. The best-performing classifier performance was reported in Table 3; the complete results are provided in

**Table 3:** The performance of machine learning-based classifiers developed using embeddings of protein language models.

| ML Models | Features | Training set |  |  |  |  | Testing set |  |  |  |  |
| --- | --- | --- | --- | --- | --- | --- | --- | --- | --- | --- | --- |
|  |  | Sn | Sp | ACC | AUC | MCC | Sn | Sp | ACC | AUC | MCC |
| MLP | ESM2_T30 | 0.83 | 0.91 | 0.87 | 0.93 | 0.75 | 0.88 | 0.88 | 0.88 | 0.94 | 0.76 |
| MLP | ESM2_T6 | 0.85 | 0.94 | 0.88 | 0.94 | 0.77 | 0.77 | 0.97 | 0.87 | 0.94 | 0.76 |
| MLP | ESM2_T12 | 0.85 | 0.91 | 0.88 | 0.94 | 0.76 | 0.65 | 0.99 | 0.82 | 0.92 | 0.69 |
| SVM | ProtBERT | 0.82 | 0.82 | 0.82 | 0.9 | 0.64 | 0.83 | 0.85 | 0.84 | 0.90 | 0.68 |
| SVM | BioBERT | 0.77 | 0.85 | 0.81 | 0.89 | 0.63 | 0.8 | 0.86 | 0.83 | 0.89 | 0.66 |

#Sn: Sensitivity, Sp: Specificity, ACC: Accuracy, AUC: Area under ROC curve, MCC: Matthews Correlation Coefficient

Supplementary Table S4. Among the evaluated embeddings, ESM2-T30 achieved the highest performance with an external AUC of 0.94 and MCC of 0.76, closely followed by ESM2-T6 (AUC of 0.94, MCC of 0.76). ESM2-T12 also showed competitive performance (AUC of 0.92, MCC of 0.69), whereas ProtBERT and BioBERT exhibited comparatively lower predictive ability, with external AUCs of 0.90 and 0.89, respectively. Based on their superior performance and ability to capture informative sequence representations, ESM2-T6 and ESM2-T30 were selected for subsequent feature-fusion and optimization analyses.

#### 3.2.3. Selection of the Best Classifier for the predictor

Since the performance of the classifiers varied across feature representations, a model-agnostic comparison of their performance was conducted. The comparison was based on average ranks calculated for each classifier using AUC. By using ranks, the performance bias related to particular features was eliminated. As shown in Figure 6 below, the CatBoost classifier achieved the best performance, with the lowest average rank (∼2.1). XGB showed the second result (∼2.7), while RF and SVM were in the middle of the ranking. Finally, LR, KNN, Naïve Bayes, and DT showed the worst performance.

**Figure 6:**
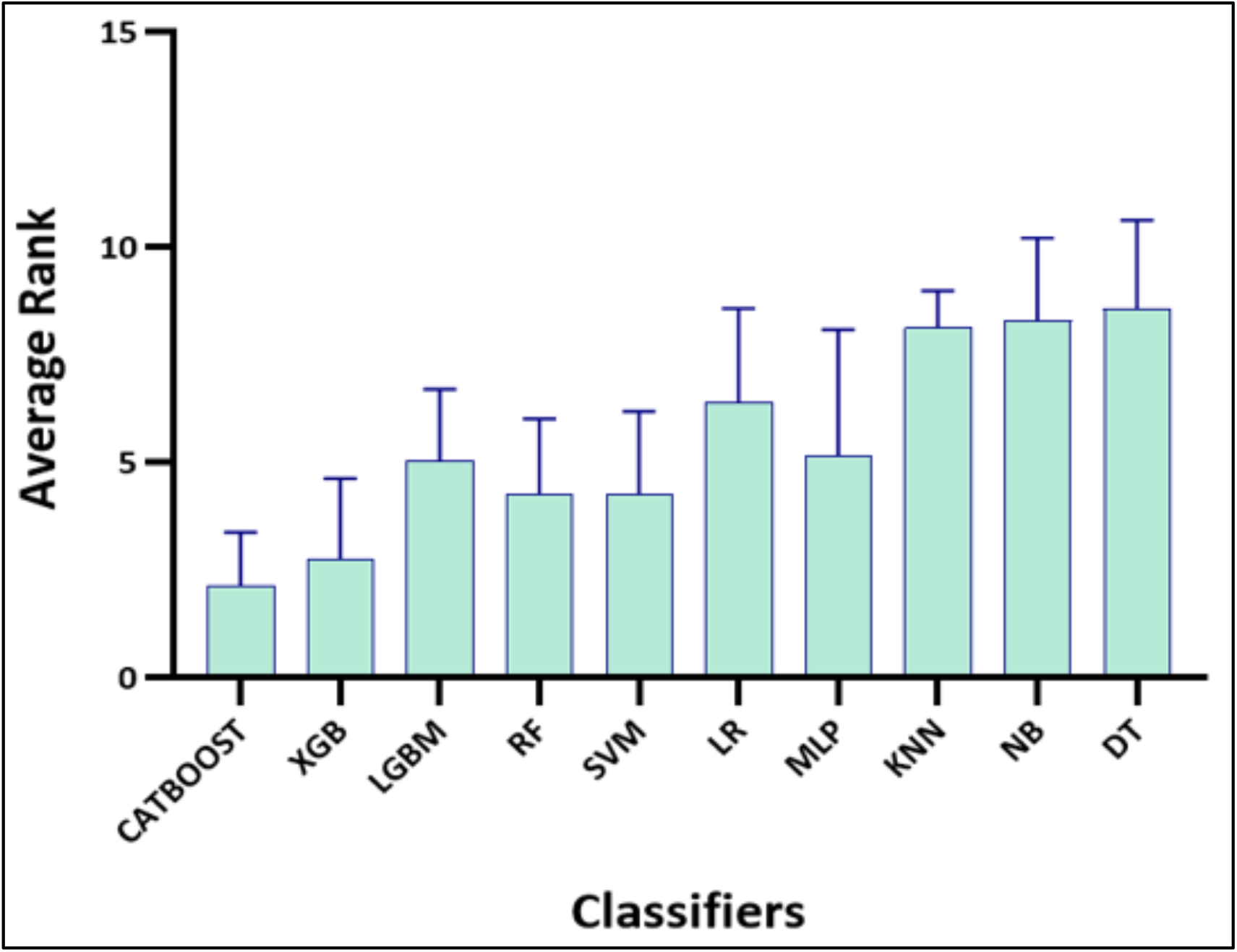
Bar plot showing the average rank-based performance of ML classifiers.

In order to determine whether the performance differences between the classifiers are statistically significant, a Friedman test was performed. The results of the Friedman test showed that the differences between all classifiers were significant Friedman = p < 0.05, p < 0.05. Toidentify which specific classifiers differ from each other, a Nemenyi post-hoc test was performed (Supplementary Figure S1). The post hoc analysis revealed that CatBoost significantly outperformed all other classifiers except XGB. Since CatBoost achieved the best average rank, it was selected as the final classifier for the fused data and selected features. It should be noted that the best-performing classifier was selected without considering specific features; thus, it may yield different results across different input representations.

### 3.3. Performance of Feature Fusions

After selecting the best classifier, we performed feature fusion using the descriptors identified during the preceding search (see previous section). Descriptors with high predictive performance (external AUC ≥ 0.93) were combined to evaluate if information from complementary sequences had a positive effect on classification performance. We tested 45 combinations using CatBoost, and the complete results are provided in Supplementary Table S5.

Among the evaluated combinations shown in Table 4, several fused feature sets outperformed their individual counterparts. The best-performing combination was OHE with the PAAC, achieving an external AUC of 0.944 and MCC of 0.80. Notably, all top-performing combinations incorporated OHE features, highlighting the importance of positional information and its complementary relationship with composition-based descriptors. Furthermore, the close agreement between training and testing performance metrics across these combinations indicates strong generalization performance and suggests that feature fusion did not introduce substantial overfitting despite the increased dimensionality. Overall, the results demonstrate that combining positional and compositional information consistently improves AOP prediction and yields a more informative representation than individual feature descriptors alone.

**Table 4:** The performance of the CatBoost model on the top 5 Pairwise Fused Feature Combinations.

| Feature fused with OHE | Training set |  |  |  |  | Testing set |  |  |  |  |
| --- | --- | --- | --- | --- | --- | --- | --- | --- | --- | --- |
|  | Sn | Sp | ACC | AUC | MCC | Sn | Sp | ACC | AUC | MCC |
| PAAC | 0.862 | 0.929 | 0.896 | 0.949 | 0.793 | 0.868 | 0.931 | 0.899 | 0.944 | 0.8 |
| SER | 0.919 | 0.914 | 0.892 | 0.946 | 0.786 | 0.944 | 0.842 | 0.893 | 0.943 | 0.79 |
| AAC | 0.863 | 0.927 | 0.895 | 0.948 | 0.792 | 0.861 | 0.931 | 0.896 | 0.942 | 0.794 |
| RRI | 0.925 | 0.919 | 0.891 | 0.944 | 0.784 | 0.944 | 0.828 | 0.886 | 0.942 | 0.777 |
| APAAC | 0.925 | 0.92 | 0.89 | 0.948 | 0.783 | 0.931 | 0.855 | 0.893 | 0.941 | 0.788 |

#Sn: Sensitivity, Sp: Specificity, ACC: Accuracy, AUC: Area under ROC curve, MCC: Matthews Correlation Coefficient, OHE: One-Hot Encoding, PAAC: Pseudo Amino Acid Composition, SER: Shannon Entropy of Residues, AAC: Amino Acid Composition, RRI: Repetitive Residue Information, APAAC: Amphiphilic Pseudo Amino Acid Composition

### 3.4. Feature Selection through SVC-L1 on top Fused Feature Representations

To further optimize the top fused feature sets, SVC-L1 was employed as a feature selection method to identify informative dimensions from the five best-performing fused representations using different regularization strengths (C = 1, 0.1, 0.01, and 0.001). The objective was to remove redundant features while preserving predictive performance. The selected feature subsets were subsequently evaluated using CatBoost, and the top results are shown in Table 5.

**Table 5:** Performance of the CatBoost Model built on the Top 6 SVC-L1-selected features across different C coefficients.

| Feature fused with OHE | C | Training set |  |  |  |  | Testing set |  |  |  |  |
| --- | --- | --- | --- | --- | --- | --- | --- | --- | --- | --- | --- |
|  |  | Sn | Sp | ACC | AUC | MCC | Sn | Sp | ACC | AUC | MCC |
| PAAC | 0.1 | 0.850 | 0.926 | 0.888 | 0.944 | 0.778 | 0.851 | 0.934 | 0.893 | 0.945 | 0.788 |
| PAAC | 1.0 | 0.863 | 0.932 | 0.897 | 0.949 | 0.797 | 0.845 | 0.944 | 0.894 | 0.944 | 0.793 |
| AAC | 1.0 | 0.859 | 0.933 | 0.896 | 0.948 | 0.795 | 0.855 | 0.934 | 0.894 | 0.944 | 0.791 |
| RR1 | 0.1 | 0.847 | 0.917 | 0.882 | 0.943 | 0.766 | 0.828 | 0.944 | 0.886 | 0.943 | 0.777 |
| SER | 1.0 | 0.858 | 0.907 | 0.883 | 0.944 | 0.767 | 0.845 | 0.931 | 0.888 | 0.943 | 0.778 |
| RRI | 1.0 | 0.855 | 0.924 | 0.889 | 0.943 | 0.781 | 0.832 | 0.944 | 0.888 | 0.941 | 0.781 |

#Sn: Sensitivity, Sp: Specificity, ACC: Accuracy, AUC: Area under ROC curve, MCC: Matthews Correlation Coefficient, OHE: One-Hot Encoding, PAAC: Pseudo Amino Acid Composition, SER: Shannon Entropy of Residues, AAC: Amino Acid Composition, RRI: Repetitive Residue Information, APAAC: Amphiphilic Pseudo Amino Acid Composition

Overall, moderate regularization (C = 1 and 0.1) substantially reduced dimensionality with minimal impact on model performance. The best result was obtained on OHE with PAAC at C = 0.1, achieving an external AUC of 0.945 with 276 selected features, slightly outperforming the fused representation without feature selection (see Table 4). Similar trends were observed for other fused feature sets. In contrast, strong regularization (C = 0.001) reduced the feature space to only 4-5 dimensions and resulted in a marked decline in predictive performance (AUC ≈ 0.85-0.87), indicating the loss of important discriminative information.

### 3.5. Alignment-Based Method - Similarity Search using BLAST

In the present study, BLASTp-short was used to perform similarity searches against the training set of AOPs and non-AOPs, with E-values ranging from 10^-4^ to 10^3^. According to the data provided in Table 6, the best results were achieved with higher E-values, reporting 102 correct hits and 11 incorrect hits for AOPs. Indeed, at lower E-values, the searches were limited to distantly similar sequences, thus reducing the coverage of the sequence space and, therefore, missing functionally relevant homologs. By increasing E-values, both sequence coverage and sensitivity were increased, allowing the identification of additional functionally related peptides and, consequently, more correct hits. Although higher E-values are usually associated with a greater likelihood of false-positive hits, the number of incorrect hits in the current study was still limited, suggesting that most additional hits are indeed functionally related.

**Table 6:** Number of correct, incorrect, and total hits for antioxidant sequences in training and testing sets for different E-values.

| E-value | Number of hits | Correct positive | Correct negatives | Incorrect hits |
| --- | --- | --- | --- | --- |
| 0.0001 | 11 | 11 | 0 | 0 |
| 0.001 | 15 | 15 | 0 | 0 |
| 0.01 | 20 | 20 | 0 | 0 |
| 0.1 | 29 | 29 | 0 | 0 |
| 1 | 46 | 39 | 6 | 1 |
| 10 | 83 | 45 | 33 | 5 |
| 20 | 95 | 46 | 41 | 8 |
| 50 | 109 | 52 | 47 | 10 |
| 100 | 113 | 52 | 50 | 11 |
| 500 | 119 | 52 | 55 | 12 |
| 1000 | 122 | 55 | 55 | 12 |

### 3.6. Ensemble Approach

To further explore potential performance gains, we investigated hybrid approaches that combine ML and similarity-based methods. Based on the preceding analyses, the best model, CatBoost with the OHE and PAAC feature set, was subsequently integrated with a BLASTp-short similarity search module optimized at an E-value threshold of 100. As shown in Figure 7, the hybrid CatBoost-BLAST framework achieved an AUC of 0.956 on the independent test dataset, indicating only a modest increase.

**Figure 7:**
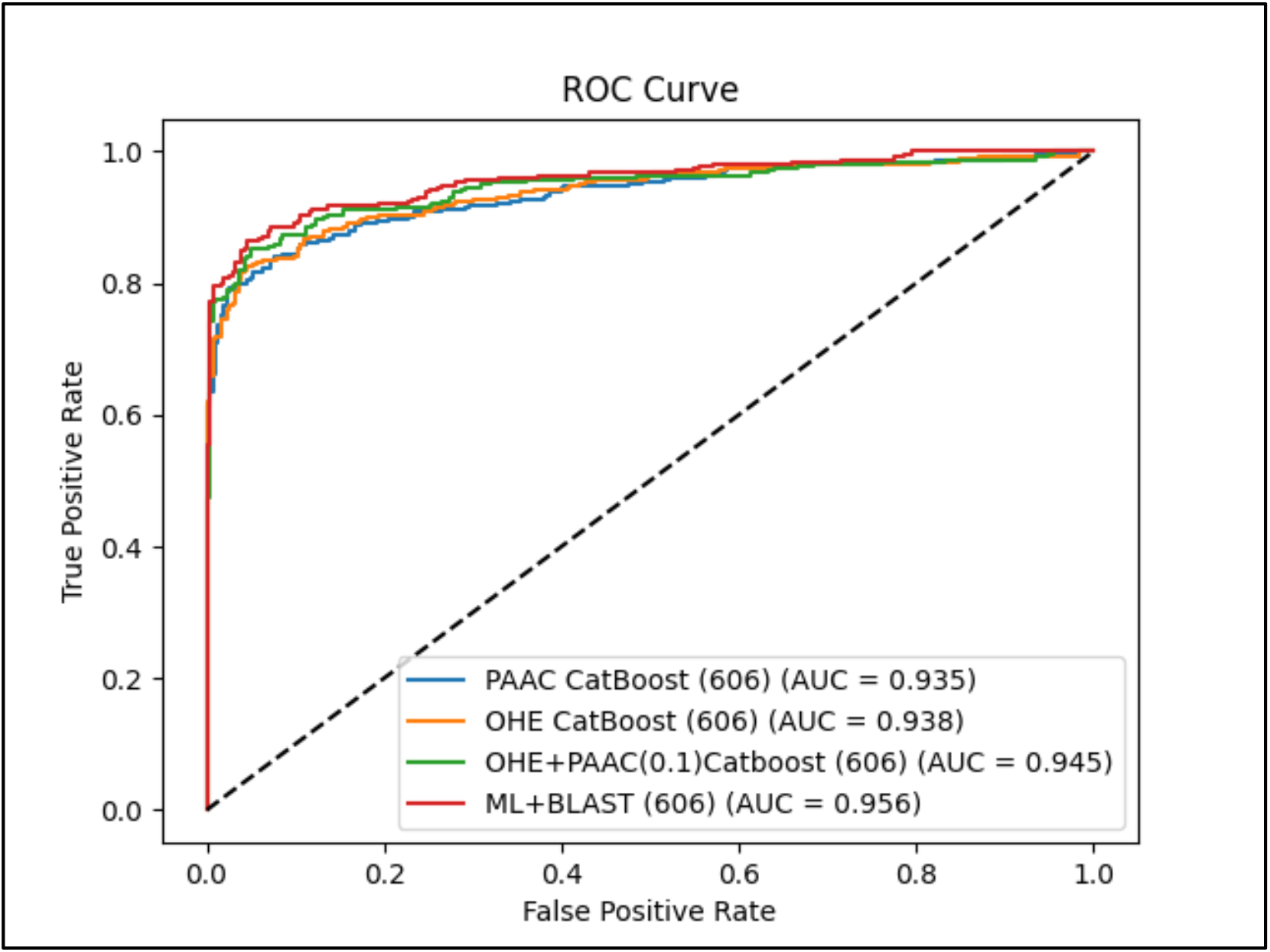
Comparison of ROC curves of the optimized CatBoost models and the hybrid framework on the testing dataset.

### 3.7. Benchmarking

To evaluate the generalizability of AntiOxiPred, benchmarking was performed on two independent datasets and compared against several state-of-the-art AOP predictors, including AOPxSVM, UniDL4BioPep, AnOxPePred, AnOxPP, AOP-DRL, and ESM4AO. Across both datasets, AntiOxiPred consistently achieved the best overall performance, demonstrating superior discrimination between AOPs and non-AOPs. On Independent Dataset 1, consisting of 150 sequences (Table 7), the AntiOxiPred ML method achieved an AUC of 0.946 and MCC of 0.841, outperforming all competing methods. Similarly, on Independent Dataset 2, comprising 250 sequences, the AntiOxiPred ML method maintained strong predictive performance, with an AUC of 0.918 and an MCC of 0.737, ranking among the best-performing predictors, as reported in Table 8. These results indicate that the model generalizes effectively across datasets originating from different sources. Although the ML+BLAST ensemble achieved the highest performance on the primary validation set, its performance varied across external benchmark datasets, likely reflecting differences in sequence similarity to the training data. In contrast, the ML model demonstrated more consistent performance across datasets.

**Table 7:** Benchmarking of AntiOxiPred against existing AOP predictors on Independent Dataset 1.

| Methods | Sn | Sp | ACC | AUC | MCC |
| --- | --- | --- | --- | --- | --- |
| AntiOxiPred | 0.893 | 0.947 | 0.92 | 0.946 | 0.841 |
| Our Hybrid (ML+BLAST) | 0.866 | 0.946 | 0.907 | 0.943 | 0.816 |
| AOPxSVM | 0.907 | 0.84 | 0.87 | 0.937 | 0.748 |
| UniDL4BioPep | 0.573 | 0.56 | 0.567 | 0.569 | 0.133 |
| AnOxPePred | 0.933 | 0.253 | 0.593 | 0.62 | 0.255 |
| AnOxPP | 0.88 | 0.907 | 0.893 | 0.942 | 0.787 |
| AOP-DRL | 0.453 | 0.587 | 0.52 | 0.579 | 0.04 |
| ESM4AO | 0.68 | 0.32 | 0.5 | 0.522133 | 0 |

#Sn: Sensitivity, Sp: Specificity, ACC: Accuracy, AUC: Area under ROC curve, MCC: Matthews Correlation Coefficient

**Table 8:** Benchmarking of AntiOxiPred against existing AOP predictors on Independent Dataset 2.

| Methods | Sn | Sp | ACC | AUC | MCC |
| --- | --- | --- | --- | --- | --- |
| AntiOxiPred | 0.848 | 0.888 | 0.868 | 0.918 | 0.737 |
| Our Hybrid (ML+BLAST) | 0.72 | 0.872 | 0.8 | 0.899 | 0.6 |
| AOPxSVM | 0.84 | 0.832 | 0.836 | 0.908 | 0.672 |
| UniDL4BioPep | 0.504 | 0.552 | 0.528 | 0.548 | 0.056 |
| AnOxPePred | 0.579 | 0.677 | 0.628 | 0.619 | 0.258 |
| AnOxPP | 0.856 | 0.88 | 0.868 | 0.906 | 0.736 |
| AOP-DRL | 0.568 | 0.6 | 0.584 | 0.636 | 0.168 |
| ESM4AO | 0.752 | 0.304 | 0.528 | 0.527 | 0.063 |

#Sn: Sensitivity, Sp: Specificity, ACC: Accuracy, AUC: Area under ROC curve, MCC: Matthews Correlation Coefficient

Therefore, the CatBoost-based model was selected as the default prediction engine, while the ML+BLAST ensemble was retained as an optional similarity-assisted prediction mode.

### 3.8. Web server implementation

To facilitate the practical application of the proposed framework, a user-friendly web server, AntiOxiPred (https://webs.iiitd.edu.in/raghava/antioxipred/), was developed to provide an interactive interface for AOP prediction. The server consists of two main modules. The “Predict” module utilizes the default CatBoost-based model and allows users to submit peptide sequences either directly or through file upload. For each sequence, the server returns the predicted class (AOPs or non-AOPs) along with the corresponding prediction score and enables users to download the results for further analysis. In addition, the hybrid model option combines an ML model with similarity-based sequence searching to offer an alternative prediction strategy. The second “BLAST Search” module identifies homologous AOPs and non-AOPs within the curated database. To promote reproducibility and accessibility, the datasets, standalone package, source code, GitHub repository (https://github.com/raghavagps/Antioxipred), and pip-installable version of AntiOxiPred (https://pypi.org/project/antioxipred/) have also been made available. Collectively, these resources provide a comprehensive platform for predicting and analyzing AOPs.

## 4. Discussion and Conclusion

Antioxidant peptides (AOPs) are becoming an important class of bioactive molecules with a wide range of potential applications in food, nutraceutical, and pharmaceutical industries. However, despite the existence of several computational predictors, their predictive accuracy is lower when tested on independent datasets.[27] We present AntiOxiPred, an ML framework that has demonstrated performance comparable to or superior to other tools on independent benchmark datasets.

The compositional analysis indicated that AOPs were enriched in L, P, Y, and A, which is consistent with previous biochemical studies.[47][48] In this context, it is notable that hydrophobic amino acids, including L and A, interact with lipid radicals, whereas Y has hydrogen-donating capacity, thereby contributing to radical scavenging. Thus, the results of the current study are consistent with previous findings, indicating that both amino acid composition and sequence structure are important determinants of AOP activity.

The optimal prediction model was selected out of ten ML algorithms evaluated using multiple sequence-derived descriptors and PLM embeddings. Among all evaluated classifiers, CatBoost consistently achieved the highest predictive performance across diverse feature representations. The superior performance of CatBoost is likely due to its ordered boosting strategy, which effectively models complex nonlinear relationships while reducing prediction bias and overfitting. Similar observations have been reported in several biological prediction studies, highlighting the effectiveness of CatBoost as a prediction model for heterogeneous biological feature representations.[49][50]

The comparison of the different feature representations showed that traditional sequence-derived descriptors still carry substantial information about AOPs. Feature fusion was beneficial for prediction, as demonstrated by the best-performing combination of OHE + PAAC, with an AUC of 0.945 and an MCC of 0.788, while the feature selection with the SVC-L1 algorithm successfully removed redundant information without deteriorating predictive power. Comparison of our method with existing AOP predictors also showed that AntiOxiPred achieves better or comparable performance, as evidenced by high AUC values on both independent sets (0.946 and 0.918). We thus conclude that our best-performing feature combination, when used with the CatBoost algorithm, is a valuable predictor of AOPs. The best standalone ML model showed consistently high performance across the independent sets, thus suggesting better extrapolation ability than the optional BLASTp-short database search module incorporated into the web server to provide additional homology-based evidence.

Although AntiOxiPred has certain advantages, the method has several limitations. The approach is currently restricted to binary classification because an extensive standard database of quantitative antioxidant data is lacking. Partly validated key events and the absence of additional experimental data on the target AOP hindered further improvement in this study. Further studies are needed to develop a regression model to predict antioxidant potency by integrating larger training sets with experimental data, incorporating structural sequence data, more powerful PLMs, and molecular information on interaction sites. Overall, AntiOxiPred provides a reliable and accurate framework for AOP prediction and can be further generalized in future studies. The web server and standalone programs offer convenient access and rapid execution for AOP screening and are available at https://webs.iiitd.edu.in/raghava/antioxipred/.

## Funding Source

The present work was supported by the Department of Biotechnology (DBT) under the grant BT/PR40158/BTIS/137/24/2021.

## Conflict of interest

No authors declare any competing financial and non-financial interests.

## Acknowledgements

We are thankful to the Council of Scientific and Industrial Research (CSIR) and the Department of Biotechnology (DBT) for providing fellowships and financial support, and to the Department of Computational Biology at IIITD, New Delhi, for the required facilities and infrastructure for the study. We also like to thank Draw.io for creating images.

## Authors’ contributions

AK manually collected, curated, and analyzed the data. AK developed the backend and frontend of the web server. AK and NB analyzed the results and finalized them with GPSR. AK, NB and GPSR prepared and reviewed the manuscript. GPSR and NB coordinated the project, which GPSR conceived. All authors read and approved the final version of the manuscript.

## Notes

### Competing Interest Statement

The authors have declared no competing interest.

